# Multimodal single cell data integration challenge: results and lessons learned

**DOI:** 10.1101/2022.04.11.487796

**Authors:** Christopher Lance, Malte D. Luecken, Daniel B. Burkhardt, Robrecht Cannoodt, Pia Rautenstrauch, Anna Laddach, Aidyn Ubingazhibov, Zhi-Jie Cao, Kaiwen Deng, Sumeer Khan, Qiao Liu, Nikolay Russkikh, Gleb Ryazantsev, Uwe Ohler, NeurIPS 2021 Multimodal data integration competition participants, Angela Oliveira Pisco, Jonathan Bloom, Smita Krishnaswamy, Fabian J. Theis

## Abstract

Biology has become a data-intensive science. Recent technological advances in single-cell genomics have enabled the measurement of multiple facets of cellular state, producing datasets with millions of single-cell observations. While these data hold great promise for understanding molecular mechanisms in health and disease, analysis challenges arising from sparsity, technical and biological variability, and high dimensionality of the data hinder the derivation of such mechanistic insights. To promote the innovation of algorithms for analysis of multimodal single-cell data, we organized a competition at NeurIPS 2021 applying the Common Task Framework to multimodal single-cell data integration. For this competition we generated the first multimodal benchmarking dataset for single-cell biology and defined three tasks in this domain: prediction of missing modalities, aligning modalities, and learning a joint representation across modalities. We further specified evaluation metrics and developed a cloud-based algorithm evaluation pipeline. Using this setup, 280 competitors submitted over 2600 proposed solutions within a 3 month period, showcasing substantial innovation especially in the modality alignment task. Here, we present the results, describe trends of well performing approaches, and discuss challenges associated with running the competition.

## 1. Introduction

Human life is possible through the function and interplay of approximately 37 trillion cells in the human body (Bianconi et al., 2013) organized into tissues, organs, and systems. The function of these systems is mediated by individual cells, the processes that occur inside them, and how they interact. As a human develops from a single fertilized egg cell, all of our cells contain the same DNA that was copied trillions of times. Yet, cells have different behaviors, functions, and proliferate at different rates. Understanding this diversity of cellular states and how they occur is the key to gaining mechanistic insight into how tissues function or malfunction in health and disease (Wagner et al., 2016).

Cellular heterogeneity can be characterized by the regulation of gene expression. Regions of DNA called genes are transcribed into messenger RNA (mRNA) molecules and subsequently translated into proteins. Various regulatory mechanisms affect gene expression, and DNA accessibility, protein abundance, and mRNA concentrations all provide complementary information on cellular state.

In the past decade, the advent of single-cell genomics technologies has enabled the measurement of these modalities in single cells (Stark et al., 2019). Single-cell RNA-sequencing measures the expression of all protein coding genes (roughly 20,000 in humans) in up to millions of cells in a single dataset (Cao et al., 2020). At a similar throughput, single-cell ATAC-sequencing measures the accessibility of DNA regions in hundreds of thousands of peaks as features (Buenrostro et al., 2015). These technologies have enabled the study of biology at an unprecedented scale and resolution, leading to detailed maps of early human embryonic development (Cao et al., 2020; Domcke et al., 2020), the discovery of new disease-associated cell types (Montoro et al., 2018), and cell-targeted therapeutic interventions (Sachs et al., 2020). Moreover, with recent advances in experimental techniques it is now possible to measure several of these modalities in the same cell (Efremova and Teichmann, 2020). While multimodal single-cell data is increasingly available, methods to analyse these data are still scarce. Due to the small volume of a single cell, measurements in any modality are sparse and noisy. Differences in molecular sampling depths between cells (sequencing depth) (Vallejos et al., 2017), and technical effects from handling cells in batches (batch effects) (Luecken et al., 2022) can often overwhelm biological differences. When analysing multimodal data, one additionally has to account for different feature spaces, as well as shared and unique variation between modalities and between batches (Argelaguet et al., 2021).

These challenges are not unique to this data type. Indeed, domain adaptation (Csurka, 2017) and multi-view learning (Zhao et al., 2017) are known problems in machine learning that are directly applicable to multimodal data integration. To promote cross-fertilization of ideas between experts in machine learning and single-cell biology and encourage method development, we organized the first competition on molecular data for NeurIPS 2021, called Multimodal Single-cell Data Integration (openproblems.bio/neurips_2021). We generated the largest, realistic benchmarking dataset currently available for multimodal single-cell data, defined three tasks and metrics to evaluate them, and developed an infrastructure for automated evaluation of user-submitted solutions (Luecken et al., 2021). Here, we briefly outline the setup of the competition, summarize the outcomes, and detail our learnings along the way.

## 2. Competition setup

We set up the Multimodal Single-cell Data Integration competition following the principles of the Common Task Framework (CTF) (Donoho, 2017) as previously described (Luecken et al., 2021). Briefly, this includes a realistic dataset that is divided into public training data and held-out private test data, a public competition, and a scoring process.

### 2.1. Dataset

Substantial effort of setting up the competition was dedicated to the generation of a bench-marking dataset that mirrors realistic challenges in multimodal single-cell data integration. In a large collaborative effort we processed bone marrow samples from 12 donors via two recent multiomic single cell technologies: CITE-seq (Stoeckius et al., 2017), which captures single-cell RNA gene expression (GEX) and surface proteins levels (as Antibody Derived Tags, ADT); and the 10x Multiome assay, which captures chromatin accessibility (based on the Assay for Transposase-Accessible Chromatin, ATAC) and single nucleus RNA gene expression (GEX) levels. After quality control, the CITE-seq and Multiome data included 90,000 and 70,000 cells, respectively. To ensure a realistic data integration problem, we distributed the data generation across 4 sites, which led to a nested batch effect structure of donor and site (Fig 4a,b). A detailed description of the data generation and analysis is available in the NeurIPS publication of the competition (Luecken et al., 2021).

### 2.2. Tasks

The CTF requires mathematically precise definitions of tasks and metrics to drive algorithm development. Here we motivate and formalize our three key multimodal tasks and related Lance et al. metrics. The full description of each task (Fig 4c-e) can be found in our previous work (Luecken et al., 2021).

#### Task 1: Predicting one modality from another

In general, genetic information flows from DNA to RNA to proteins. DNA must be accessible (ATAC data) to produce RNA (GEX data), and RNA in turn is used as a template to produce proteins (ADT data). The goal of this task is to accurately predict one modality from another with performance measured using root mean squared error. Algorithms capable of performing well at this task may learn rules governing the complex regulatory interactions between layers of genetic information.

#### Task 2: Matching cells between modalities

Most existing single-cell datasets measure a single modality. Aligning modalities while preserving underlying biology will enable leveraging complementary layers of information measured independently. The goal of this task is to correctly match multimodal profiles measured from the same cell when the correspondences are hidden. Performance is based on the weight put on the correct pairings, capturing the probability that a cell is correctly matched. Understanding how feature selection influences matching accuracy may shed light on the significance of different regions of DNA or transcripts of RNA in cell identity and regulation of downstream genetic processes.

#### Task 3: Jointly learning representations of cellular identity

Multimodal measurements combine complementary layers of information. The goal of this tasks is to learn highly resolved but lower-dimensional representations (≦ 100 dimensions) of the underlying biological states of cells while removing batch effects across sites. Performance is measured taking the average of 6 metrics assessing the degree of batch removal and conservation of biological variation detailed in Luecken et al. (2021).

## 3. Competition results

Full details of all analyses and implementations of all methods described in the following sections are organized within the Open Problems github organization https://github.com/openproblems-bio/in_repositories_neurIPS21_comp_paper_reproducibility (soon to be published) and neurips2021_multimodal_topmethods. Here we summarize the main features.

### 3.1. General trends and feedback

More than 280 participants comprising 172 teams submitted over 2600 models to our competition. We later surveyed competitors (35 teams) to learn more about participant background, the diversity of methods, and the experience of the competition.

#### Toolkit breakdown

Several toolkits exist for loading and processing single-cell data. As depicted in Figure 5, the most popular toolkit among competitors was Scanpy (46 %). Surprisingly, one third of responding teams did not use any toolkit. Among deep learning frameworks, PyTorch was twice as popular as TensorFlow despite Tensorflow having approximately 3 times as many monthly downloads from PyPI compared to PyTorch (https://pypistats.org). This matches the previously reported trend of increasing popularity of PyTorch in research-oriented projects (He, 2019).

#### General feedback

To learn more about highlights and issues participants experienced during the competition, we asked for their main positive and challenging aspects.

Due to supply chain issues and barriers in data generation, we had to change deadlines and data delivery dates through the competition. This was the source of the main criticism we received. As a lesson learned, especially for data of such kind, a large time buffer for the data generation process is required. Changing dates and deadlines should be avoided.

The second most common issue raised by participants was related to the evaluation infrastructure. The turnaround time to get the evaluation results frustrated competitors, with some not knowing how final submissions performed until after the deadline. Therefore, one should anticipate a surge in demand for evaluation resources during the final phase of a competition and carefully assess the computational demands.

60 % of respondents planned to make further use of our benchmarking dataset, highlighting the value of a purpose-built benchmarking dataset. We also received feedback that lack of availability of the full dataset at competition start frustrated attempts to participate. However, the data provided as a cleaned up .h5ad file prepared by the censoring components in our starter kits was often mentioned as a great way to get started with method development quickly.

### 3.2. Task 1: Modality Prediction

#### 3.2.1. Overview of the modality prediction competition

The modality prediction task was the most accessible to a non-biological audience, drawing 1,393 submissions. This was expected due to Task 1’s resemblance to a standard machine translation or prediction task. This popularity suggests the importance of including a straightforward task setup when introducing ML practitioners to a new data domain.

Task 1 contained 5 subtasks: predicting one modality from the other in each direction for each technology (CITE-seq: ADT2GEX, GEX2ADT and Multiome: ATAC2GEX, GEX2ATAC) and the best average performance. In general, most submitted solutions performed similarly (Fig 1 a). While GEX2ATAC and ADT2GEX submissions showed little improvement over baseline, some ATAC2GEX and GEX2ADT submissions outperformed the baseline and most other submissions. In these subtasks, survey results showed that most top ranked methods were deep learning models but no specific model architecture consistently outperformed others (Fig 1 b). These models typically had a depth of 3-5 hidden layers and in some cases 5-10, with several participants reporting a drop in performance with additional layers. While shallow deep learning models appear well suited to ATAC2GEX and GEX2ADT, the top GEX2ADT method used kernel ridge regression (see Section 3.2.2).

**Figure 1:**
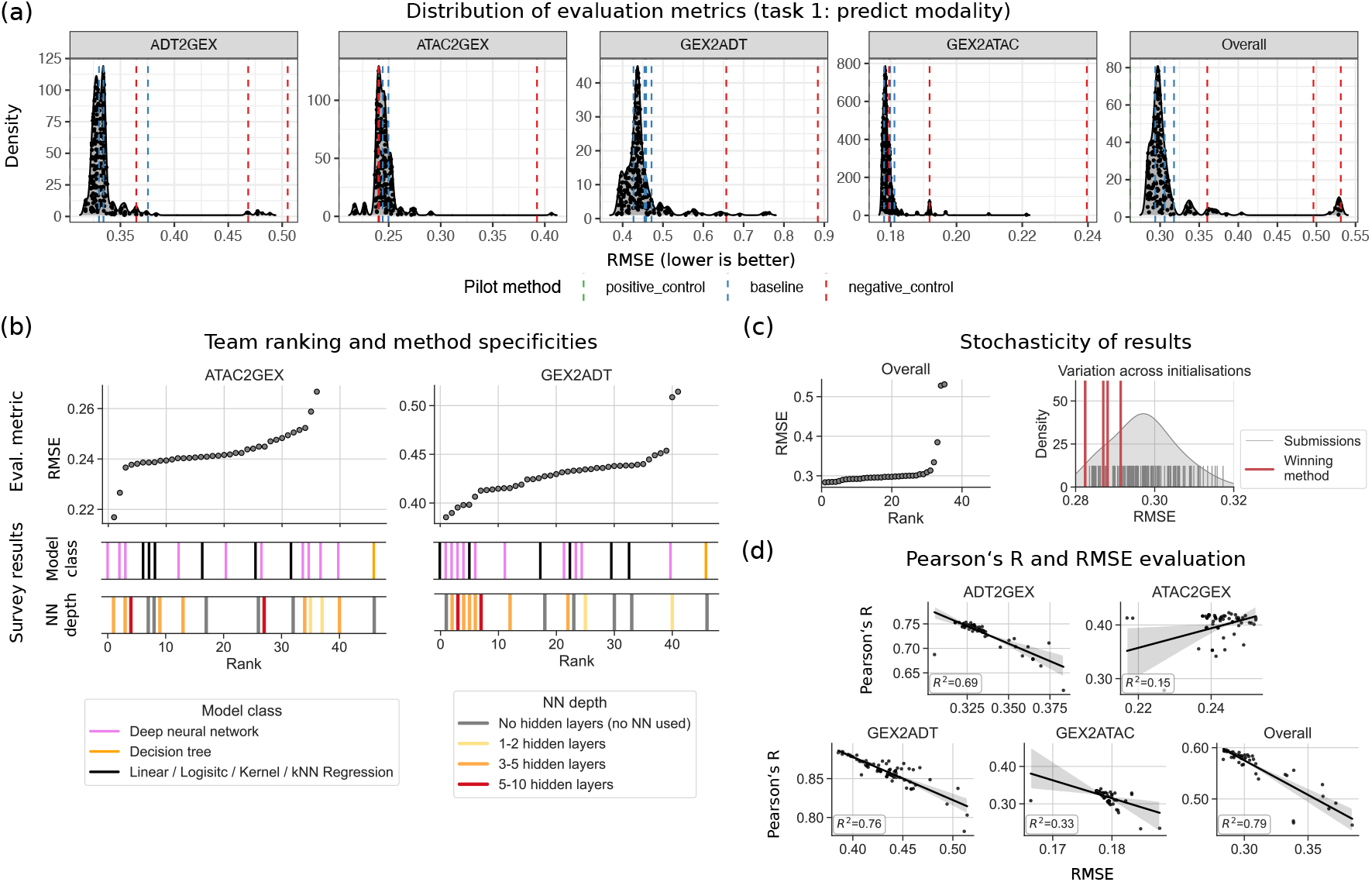
Modality prediction task results. **(a)** RMSE distribution of the evaluation metrics for the different predictions and overall summary. **(b)** Team ranking and method characteristics for ATAC2GEX and GEX2ADT. **(c)** Comparison of all submissions with reruns of the wining method. **(d)** Person’s R and RMSE evaluation of the top performing methods per task.

Further analysis of metric outputs showed that, especially in the GEX2ATAC, ADT2GEX and Overall subtasks, similar RMSE scores achieved by most submissions led non-robust rankings (Fig 1 c, left). Indeed, stochasticity in model training had a substantial effect on ranks: rerunning the Overall subtask winner (DANCE) four times led to a shift in up to 8 ranks (Fig 1 c, right). Feedback from the community and further analysis revealed two issues that likely contributed to overall similar and poor performance. First, with highly sparse and binarized ATAC data, predicting mostly zeros is effective for RMSE (see high performance of negative control in Fig 1 a. Up-weighting correctly predicted open chromatin regions may result in a more meaningful assessment. Second, scales of gene expression data differed by site, as a direct consequence of our per-sample data processing strategy (see Luecken et al. (2021) for details). This limited the generalizability of predictions to test data from an unseen site. As a straightforward solution, one could fix a relative scale for expression values. Alternatively, a correlation-based metric could be used, as suggested by several participants. In pilot phase testing on a subset of the data, we found that RMSE and correlation scores returned consistent method rankings and opted for RMSE as a more straightforward cost function to optimize. Comparing RMSE and Pearson’s R values (Fig 1 d) shows that this is still true in several subtasks. However, especially for the ATAC2GEX subtask, predicting the correct pattern may be more important than the exact value. In future competitions, using a correlation-based metric may improve overall performance.

#### 3.2.2. Selected winning methods

Here we briefly describe a subset of the winning methods that improved substantially upon the baseline. For all full method descriptions, please refer to Appendix C.1.

##### ATAC→GEX: Cajal

Team Cajal used a deep neural network consisting of a dropout layer, 4 hidden layers with ReLU activation, an output layer with no activation function (linear regression) and a final Lambda layer that clipped output to biologically reasonable values (the range present in the training data). Cell-type specific features were selected as input based on differential accessibility (t-test on binarized data) on an annotated reference dataset. Total counts, median total counts per batch, and standard deviation of total counts per batch were also used as input features after being normalized. Implementation was with TensorFlow (Abadi et al., 2015); KerasTuner (O’Malley et al., 2019) was used to optimize hyperparameters (percentage dropout, number of hidden layers, number of nodes per layer).

##### GEX→ADT: Guanlab - dengkw

Team Guanlab - dengkw used kernel ridge regression (KRR) with RBF kernel 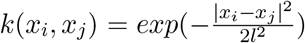. Input data was processed by applying truncated SVD to the concatenation of the training and test data from modality 1 followed by row-wise z-score normalization on the reduced matrix. The KRR model was trained to predict the truncated SVD of modality 2 from the normalized training matrix of modality 1. Predictions on the normalized test matrix were re-mapped to the modality 2 feature space via the right singular vectors. The final submission used a training strategy of random sampling and assembling to overcome memory limitations. First, they shuffled the training batches and trained two models on the first and second half of batches. They repeated this step 5 times to fit 10 models, whose 10 outputs were averaged to generate the final predictions. The number of component for modality 1 (300), modality 2 (70), length scale *l* (10), and regularization strength *α* (0.2) were optimized by cross-validation.

### 3.3. Task 2: Match Modality

#### 3.3.1. Overview of the match modality competition

As in task 1, the match modality task asked participants to re-identify the correspondence of cellular profiles across modalities in 5 subtasks: GEX2ATAC, ATAC2GEX, ADT2GEX, GEX2ADT, and Overall. Unlike task 1, one method emerged as a clear winner across all subtasks, with several others clearly distinguished from the pack (Fig 2 a). This was the least popular task with 462 submissions. Survey results show that top-ranked teams had previous experience with single-cell data analysis and used popular single-cell analysis tool kits. Moreover, top performers applied shallow deep learning models (Fig 2 c).

**Figure 2:**
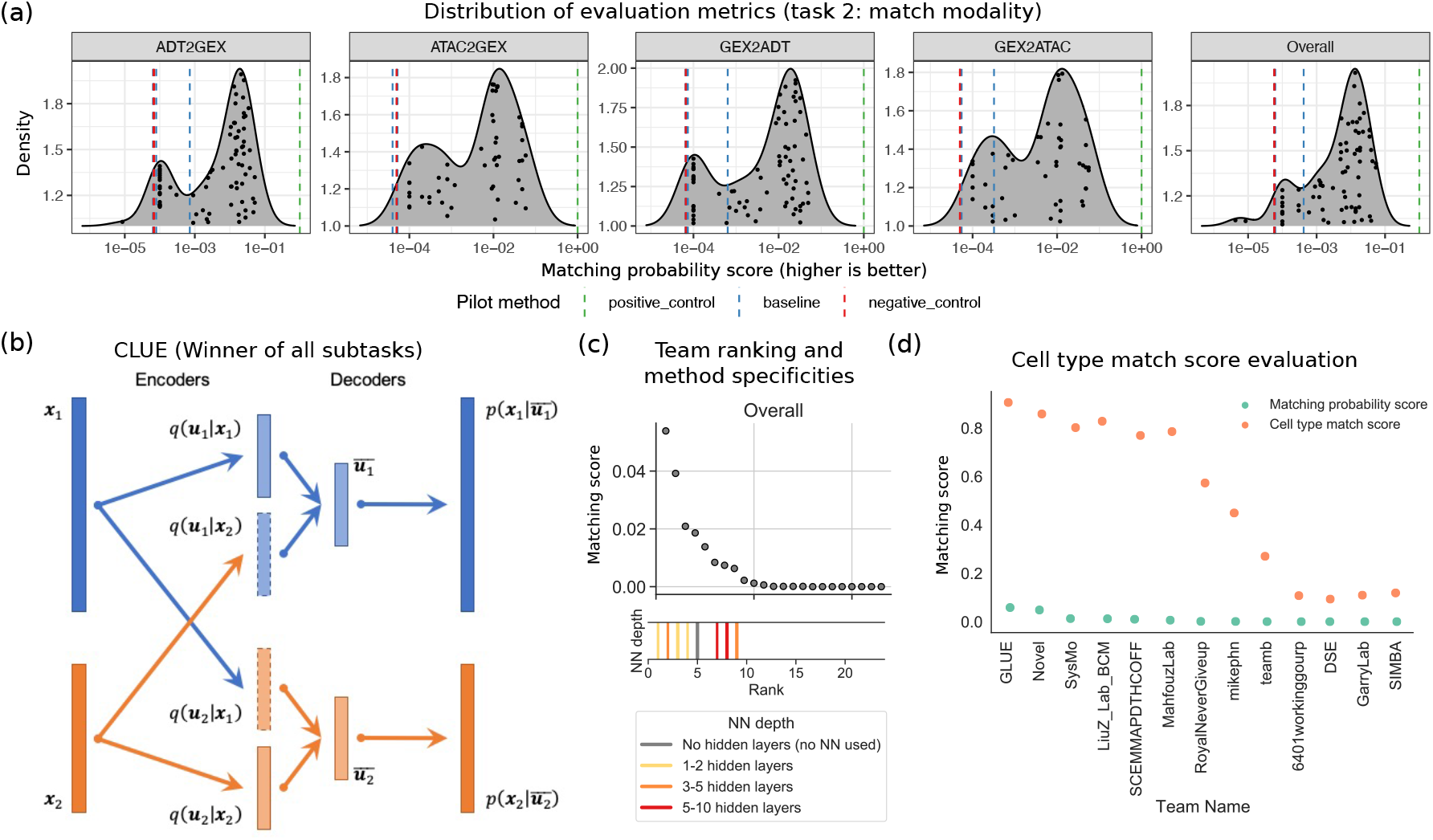
Match modality task results. **(a)** RMSE distribution of the evaluation metrics for the different predictions and overall summary. **(b)** Schematic representation of CLUE, the winner method across all sub-tasks. **(c)** Team ranking vs scores and model characteristics. **(d)** ATAC2GEX matching scores at the cell and cell-type level by team.

We expected this to task to be especially difficult due to the biological limits of identifiability. Indeed, cells of the same type are often treated as biological replicates for statistical tests, with differences ascribed to stochastic variation. When assessing matching scores based on the easier task of matching among 22 cell-type labels in the Multiome data, scores are vastly improved (Fig 2 d), with drastic separation between competitors and a top score around 0.90. In real scenarios with truly unpaired data, matching cell-type profiles with this degree of accuracy would already be impactful. Comparing the performance of top methods at the cell-type and cell level (Fig 2 c, d), we see that the top two methods have similar performance to the next four at the cell-type level, but 2-5x higher performance at the cell level. Excitingly, this suggests the top two methods are exploiting structure beyond known cell-type annotations, with the top probability matching score in the ATAC2GEX subtask (0.058 by CLUE) about 1000x higher than the expected score of a random matching of the 20,009 cells. CLUE scored between 0.049 and 0.058 on all four subtasks, indicating that the direction of matching did not significantly impact performance. However, since CITE had lower scores and fewer cells (15,066), we conclude that matching was an easier task for CLUE on Multiome than CITE. Remarkably, neither of the top two methods incorporated any knowledge of the relationship between feature sets (e.g., the distance from peaks to genes, or the correspondence of genes and proteins). This may be an avenue for further improvement.

#### 3.3.2. The winning method

##### Winner in all categories: CLUE

CLUE (Cross-Linked Universal Embedding) is a method designed for semi-supervised modality matching of single-cell multiomics data. It builds upon VAE (variational autoencoders) to learn joint cell embeddings from different modalities. Given that cell type resolution often differs across modalities, indiscriminately modeling all modalities using a single embedding space could cause information in higher-resolution modalities to be contaminated by lower-resolution ones. To overcome this issue, CLUE partitions the embedding space into modality-specific subspaces, each tasked to reconstruct one specific modality only. A matrix of encoders is then used to project each cell into all modality-specific subspaces, regardless of which modality the input cell originates from (Fig 2 b). The joint CLUE embedding is constructed by concatenating all the modality-specific subspaces. Such a design enables CLUE to preserve both shared and modality-specific information, significantly boosting the accuracy of modality matching.

### 3.4. Task 3: Joint Embedding

#### 3.4.1. Overview of the joint embedding competition

Learning a low-dimensional representation of cells from single-cell data is a common step in a typical analysis pipeline (Luecken and Theis, 2019). With the rise of multimodal data, the ability to meaningfully embed these data will be crucial for analysis and visualization. This motivation is reflected by the high number of submissions for this task (756 in total) and the wide use of single-cell analysis tool kits among participating teams (8/10 survey responses). We divided the task into pre-trained models and models that were trained online (using only test data), in order to separately evaluate models that use additional information. However, top performers scored similarly across training regimes. Interestingly, the MNN baseline method (Haghverdi et al., 2018) also performed well, resulting in only 14 out of 25 teams outperforming the baseline (Fig 3 a). As in the previous tasks, higher performing submissions tended to use shallow neural networks (Fig 3 b). Yet, the winning team of the online CITE-seq integration used a simple linear-dimensionality reduction method without explicit batch correction.

**Figure 3:**
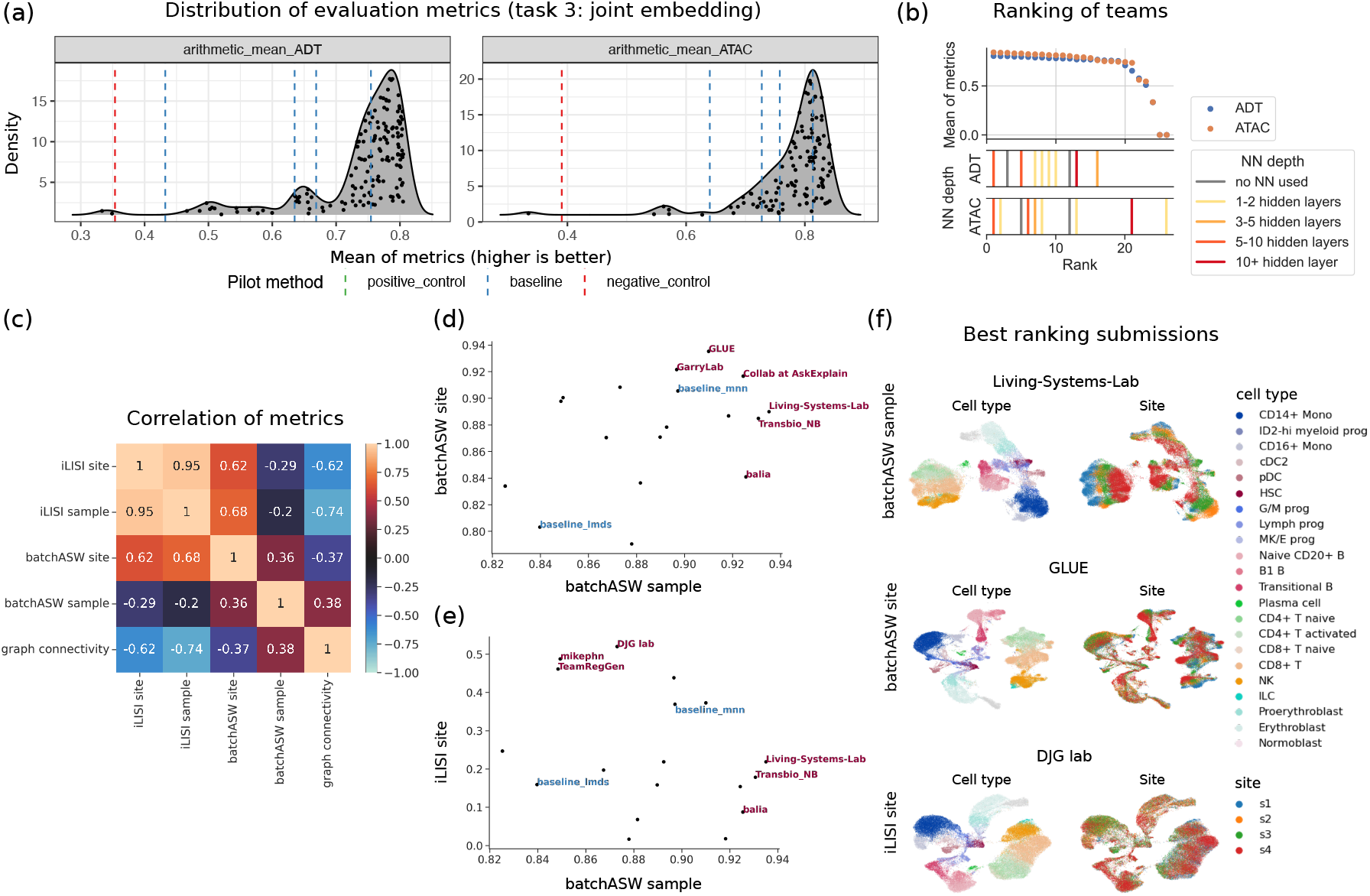
Joint Embedding task results: **(a)** Mean metric distrubtions for all submissions for the CITE-seq and Multiome tasks. **(b)** Team ranking vs method specificities. **(c)** Heatmap of Spearman rank coefficients between batch removal metric scores across all top submissions per team that were better than the LMDS-based baseline (online and pre-trained). **(d)** and **(e)** Scatterplots of the batchASW sample metric scores vs metrics accounting for site-specific batch effects; baseline models highlighted in blue, top ranking methods in red. **(f)** UMAP plots of embeddings from the top models by different batch removal metrics colored by cell type and site.

As explicit batch correction was also lacking in other winning methods, we re-evaluated our batch removal metrics. We observed that the batchASW and graph connectivity metrics applied in the challenge did not sufficiently discriminate between submissions (Fig 6). Due to computational limitations, the metrics chosen in the competition represent only a subset of batch removal metrics available (Luecken et al., 2022). Yet, the chosen metrics are either easily fulfilled (graph connectivity) or do not sufficiently account for the nested batch effect structure in the data (batchASW - based on average silhouette width). By design, our dataset contains both within-site donor variation and between-site technical variation, where the former is typically considerably smaller than the latter. However, the silhouette score only considers the nearest class cluster, which can lead to between-site differences being overlooked. As only data from a single site was available when the setup was evaluated, this effect went unnoticed.

Extending our evaluation to different batch removal metrics that can specifically account for the site batch effects, we find diverging batch removal assessments across the best-performing submissions for the Multiome dataset (Fig 3 c-e). In particular, specifically accounting for site batch effects in the batchASW metric (using *site* instead of *sample* as batch covariate) leads to higher correlations with an orthogonal metric for batch removal that accounts for all batches simultaneously, graph iLISI (integration Local Inverse Simpson’s Index (Luecken et al., 2022; Korsunsky et al., 2019)). Indeed, visualizing the embeddings of the highest-ranking submissions by each of the three batch removal metrics (Fig 3 d-f and Fig 7) highlights that batchASW site and iLISI site better capture site batch effect removal. Explicit evaluation of nested batch effects will need to be accounted for in future competitions.

#### 3.4.2. Selected winning method

Here we briefly describe one of the winning methods. For full method descriptions, please refer to Appendix C.3.

##### Multiome, pre-trained and CITE, pre-trained Amateur

Team Amateur’s method “joint embedding with a regularized autoencoder” (JAE) was inspired from previous work on scDEC (Liu et al., 2021), which aims at simultaneous deep generative modeling and clustering of single-cell data. Here, the scDEC model was simplified by removing the discriminator networks and adding constraints to the encoder latent space so that JAE requires latent features to recover more biological knowledge, including cell type, batch, and cell-cycle phase.

In the JAE model, each modality (except ADT) is first SVD transformed (e.g. to 100 components) and concatenated. The information from cell annotations (e.g., cell label, cell cycle score, and cell batch) is incorporated to constrain the structure of latent features. In this manner some latent features should recover the cell type information and some should recover the cell cycle score. Batch-related features should recover batch labels as randomly as possible to potentially eliminate this effect. Some features in the latent space were left without constraint to ensure the flexibility of network. JAE was pre-trained using the provided annotated datasets in an end-to-end fashion where multiple loss functions were used, including autoencoder reconstruction loss, cell type prediction cross entropy loss, cell cycle phase score mean squared error (MSE) loss, and batch loss. These losses were balanced, resulting in a total loss of 0.7 AE + 0.2 CT + 0.05 CC + 0.05 Batch.

## 4. Conclusion and outlook

For the first NeurIPS competition on molecular data, we generated a novel fit-for-purpose benchmarking dataset, defined three tasks with metrics to evaluate performance, and developed infrastructure to facilitate method evaluation and re-use. The 280 participants, many of whom did not have any single-cell expertise, submitted over 2600 solutions. The competition supported substantial innovation, especially on the task of matching cellular profiles across modalities.

Evaluating participant feedback and submission trends, we find that neural networks are most popular, and that top methods found relatively shallow architectures to be optimal for single-cell multimodal data. We also identified areas for technical improvements on performance metrics and task definitions. In particular, alternative metrics for modality prediction and batch correction would have better directed method development.

The infrastructure we developed to support the competition enables solutions for multimodal data integration tasks to be ported to a living benchmarking framework, which we developed in the Open Problems for Single-Cell Analysis project (https://openproblems.bio). In this context, we see the competition at NeurIPS 2021 as the start of a long-term benchmarking effort for multimodal data integration methods. Here, suites of metrics may be used to probe and rank methods, leveraging our dedicated effort to generate and share benchmarking data. Through Open Problems and competitions that engage the wider community, we hope to plant seeds that lead to computational innovations driving discovery in single-cell genomics.

## Acknowledgments

Thanks to all participants, organizers, and people generating and analyzing the data. We thank Cellarity, Saturncloud, and Biolegend for sponsorship of the competition. This project has been made possible in part by grant number 2021-235155 from the Chan Zuckerberg Initiative DAF, an advised fund of Silicon Valley Community Foundation and by the Helmholtz Association’s Initiative and Networking Fund through Helmholtz AI [ZT-I-PF-5-01] and sparse2big [ZT-I-0007]. CL is supported by the joint research school Munich School for Data Science (MUDS). PR and UO acknowledge support by DFG Research Unit FOR 2841 and the DFG International Research Training Group IRTG 2403.

## References

[1] Martffn Abadi, Ashish Agarwal, Paul Barham, Eugene Brevdo, Zhifeng Chen, Craig Citro, Greg S Corrado, Andy Davis, Jeffrey Dean, Matthieu Devin, et al. Tensorflow: Large-scale machine learning on heterogeneous systems, 2015.

[2] Martffn Abadi, Ashish Agarwal, Paul Barham, Eugene Brevdo, Zhifeng Chen, Craig Citro, Greg S Corrado, Andy Davis, Jeffrey Dean, Matthieu Devin, et al. Tensorflow: Large-scale machine learning on heterogeneous distributed systems. arXiv preprint 1603.04467, 2016.

[3] Ricard Argelaguet, Anna S. E. Cuomo, Oliver Stegle, and John C. Marioni. Computational principles and challenges in single-cell data integration. Nature Biotechnology, 39:1202–1215, may 2021. ISSN 1087-0156. doi: 10.1038/s41587-021-00895-7. URL http://www.nature.com/articles/s41587-021-00895-7.

[4] Tal Ashuach, Mariano I Gabitto, Michael I Jordan, and Nir Yosef. Multivi: deep generative model for the integration of multi-modal data. bioRxiv, 2021.

[5] Peter W Battaglia, Jessica B Hamrick, Victor Bapst, Alvaro Sanchez-Gonzalez, Vinicius Zambaldi, Mateusz Malinowski, Andrea Tacchetti, David Raposo, Adam Santoro, Ryan Faulkner, et al. Relational inductive biases, deep learning, and graph networks. arXiv preprint 1806.01261, 2018.

[6] Eva Bianconi, Allison Piovesan, Federica Facchin, Alina Beraudi, Raffaella Casadei, Flavia Frabetti, Lorenza Vitale, Maria Chiara Pelleri, Simone Tassani, Francesco Piva, Soledad Perez-Amodio, Pierluigi Strippoli, and Silvia Canaider. An estimation of the number of cells in the human body. Annals of Human Biology, 40(6):463–471, 2013. doi: 10.3109/03014460.2013.807878.URL https://doi.org/10.3109/03014460.2013.807878. PMID: 23829164.

[7] Jason D. Buenrostro, Beijing Wu, Ulrike M. Litzenburger, Dave Ruff, Michael L. Gonzales, Michael P. Snyder, Howard Y. Chang, and William J. Greenleaf. Single-cell chromatin accessibility reveals principles of regulatory variation - Nature. Nature, 523(7561):486–490, Jul 2015. ISSN 1476-4687. doi: 10.1038/nature14590.

[8] Junyue Cao, Diana R. O’Day, Hannah A. Pliner, Paul D. Kingsley, Mei Deng, Riza M. Daza, Michael A. Zager, Kimberly A. Aldinger, Ronnie Blecher-Gonen, Fan Zhang, Malte Spielmann, James Palis, Dan Doherty, Frank J. Steemers, Ian A. Glass, Cole Trapnell, and Jay Shendur∼e. A human cell atlas of fetal gene expression. Science, 370(6518): eaba7721, nov 2020. ISSN 0036-8075. doi: 10.1126/science.aba7721. URL https://www.sciencemag.org/lookup/doi/10.1126/science.aba7721.

[9] Gabriela Csurka. A Comprehensive Survey on Domain Adaptation for Visual Applications. Advances in Computer Vision and Pattern Recognition, (9783319583464):1–35, 2017. ISSN 21916594. doi: 10.1007/978-3-319-58347-1_1. URL https://link.springer.com/chapter/10.1007/978-3-319-58347-1_1.

[10] Silvia Domcke, Andrew J. Hill, Riza M. Daza, Junyue Cao, Diana R. O’Day, Hannah A. Pliner, Kimberly A. Aldinger, Dmitry Pokholok, Fan Zhang, Jennifer H. Milbank, Michael A. Zager, Ian A. Glass, Frank J. Steemers, Dan Doherty, Cole Trapnell, Darren A. Cusanovich, and Jay Shendure. A human cell atlas of fetal chromatin accessibility. Science, 370(6518):eaba7612, nov 2020. ISSN 0036-8075. doi: 10.1126/science.aba7612. URL https://www.sciencemag.org/lookup/doi/10.1126/science.aba7612.

[11] David Donoho. 50 years of data science. Journal of Computational and Graphical Statistics, 26(4):745–766, 2017. doi: 10.1080/10618600.2017.1384734. URL https://doi.org/10.1080/10618600.2017.1384734.

[12] Mirjana Efremova and Sarah A. Teichmann. Computational methods for single-cell omics across modalities. Nature Methods, 17(1):14–17, jan 2020. ISSN 1548-7091. doi: 10.1038/s41592-019-0692-4. URL http://www.nature.com/articles/s41592-019-0692-4.

[13] Laleh Haghverdi, Aaron T. L. Lun, Michael D. Morgan, and John C. Marioni. Batch effects in single-cell RNA-sequencing data are corrected by matching mutual nearest neighbors. Nature Biotechnology, 36(5):421–427, apr 2018. ISSN 1087-0156. doi: 10.1038/nbt.4091. URL http://www.nature.com/doifinder/10.1038/nbt.4091.

[14] Horace He. The state of machine learning frameworks in 2019. The Gradient, 2019.

[15] Thomas N Kipf and Max Welling. Semi-supervised classification with graph convolutional networks. arXiv preprint 1609.02907, 2016.

[16] Ilya Korsunsky, Nghia Millard, Jean Fan, Kamil Slowikowski, Fan Zhang, Kevin Wei, Yuriy Baglaenko, Michael Brenner, Po-ru Loh, and Soumya Raychaudhuri. Fast, sensitive and accurate integration of single-cell data with Harmony. Nature Methods, 16(12):1289–1296, dec 2019. ISSN 1548-7091. doi: 10.1038/s41592-019-0619-0. URL http://dx.doi.org/10.1038/s41592-019-0619-0.

[17] Qiao Liu, Shengquan Chen, Rui Jiang, and Wing Hung Wong. Simultaneous deep generative modelling and clustering of single-cell genomic data. Nature machine intelligence, 3(6): 536–544, 2021.

[18] Malte D Luecken and Fabian J Theis. Current best practices in single-cell rna-seq analysis: a tutorial. Molecular systems biology, 15(6):e8746, 2019.

[19] Malte D. Luecken, Daniel B. Burkhardt, Robrecht Cannoodt, Christopher Lance, Aditi Agrawal, Hananeh Aliee, Ann T. Chen, Louise Deconinck, Alejandro Granados, Shelly Huynh, Laura Isacco, Yang Joon Kim, Bony De Kumar, Sunil Kuppasani, Heiko Lickert, Ãaron McGeever, Joaquin Caceres Melgarejo, Maurizio Morri, Michaela F. Mueller, Bastian Rieck, Kaylie Schneider, Scott Steelman, Dan J. Treacy, Alexander Tong, Michael Sterr, Alexandra-Chloé Villani, Guilin Wang, Ce Zhang, Angela O. Pisco, Smita Krishnaswamy, Fabian J̃. Theis, and Jonathan M. Bloom. A sandbox for prediction and integration of DNA, RNA, and protein data in single cells. Technical report, 2021. URL https://openreview.net/forum?id=gN35BGa1Rthttps://openproblems.bio/neurips.

[20] Malte D. Luecken, M. Büttner, K. Chaichoompu, A. Danese, M. Interlandi, M. F. Mueller, D. C. Strobl, L. Zappia, M. Dugas, M. Colomé-Tatché, and Fabian J. Theis. Benchmarking atlas-level data integration in single-cell genomics. Nature Methods, 19(1):41–50, 2022.

[21] Daniel T. Montoro, Adam L. Haber, Moshe Biton, Vladimir Vinarsky, Brian Lin, Susan E. Birket, Feng Yuan, Sijia Chen, Hui Min Leung, Jorge Villoria, Noga Rogel, Grace Burgin, Alexander M. Tsankov, Avinash Waghray, Michal Slyper, Julia Waldman, Danielle Nguyen, Lan and D̃ionne, Orit Rozenblatt-Rosen, Purushothama Rao Tata, Hongmei Mou, Manjunatha Shivaraju, Hermann Bihler, Martin Mense, Guillermo J. Tearney, Steven M. Rowe, John F. Engelhardt, Aviv Regev, and Jayaraj Rajagopal. A revised airway epithelial hierarchy includes CFTR-expressing ionocytes. Nature, 560 (7718):319–324, aug 2018. ISSN 14764687. doi: 10.1038/s41586-018-0393-7. URL https://doi.org/10.1038/s41586-018-0393-7.

[22] Tom O’Malley, Elie Bursztein, James Long, François Chollet, Haifeng Jin, Luca Invernizzi, et al. Keras Tuner. https://github.com/keras-team/keras-tuner, 2019.

[23] Tom O’Malley, Elie Bursztein, James Long, François Chollet, Haifeng Jin, Luca Invernizzi, et al. Keras tuner. Retrieved May, 21:2020, 2019.

[24] Alec Radford, Jong Wook Kim, Chris Hallacy, Aditya Ramesh, Gabriel Goh, Sandhini Agarwal, Girish Sastry, Amanda Askell, Pamela Mishkin, Jack Clark, et al. Learning transferable visual models from natural language supervision. In International Conference on Machine Learning, pages 8748–8763. PMLR, 2021.

[25] Stephan Sachs, Aimée Bastidas-Ponce, Sophie Tritschler, Mostafa Bakhti, Anika Böttcher, Miguel A. Sánchez-Garrido, Marta Tarquis-Medina, Maximilian Kleinert, Katrin Fischer, Sigrid Jall, Alexandra Harger, Erik Bader, Sara Roscioni, Annette Ussar, Siegfried and F̃euchtinger, Burcak Yesildag, Aparna Neelakandhan, Christine B. Jensen, Marion Cornu, Bin Yang, Brian Finan, Richard D. DiMarchi, Matthias H. Tschöp, Fabian J. NeurIPS 2021 - Multimodal single cell data integration challenge

[26] Theis Susanna M. Hofmann, Timo D. Müller, and Heiko Lickert. Targeted pharmacological therapy restores $-cell function for diabetes remission. Nature Metabolism, 2 (2):192–209, feb 2020. ISSN 25225812. doi: 10.1038/s42255-020-0171-3. URL https://doi.org/10.1038/s42255-020-0171-3.

[27] Rory Stark, Marta Grzelak, and James Hadfield. RNA sequencing: the teenage years. Nature Reviews Genetics, pages 1–26, jul 2019. ISSN 1471-0056. doi: 10.1038/s41576-019-0150-2. URL http://www.nature.com/articles/s41576-019-0150-2.

[28] Marlon Stoeckius, Christoph Hafemeister, William Stephenson, Brian Houck-Loomis, Pratip K. Chattopadhyay, Harold Swerdlow, Rahul Satija, and Peter Smibert. Simultaneous epitope and transcriptome measurement in single cells. Nature Methods, 14(9):865–868, jul 2017. ISSN 1548-7091. doi: 10.1038/nmeth.4380. URL http://www.nature.com/doifinder/10.1038/nmeth.4380.

[29] Catalina A. Vallejos, Davide Risso, Antonio Scialdone, Sandrine Dudoit, and John C. Marioni. Normalizing single-cell RNA sequencing data: challenges and opportunities. Nature Methods, 14(6):565–571, may 2017. ISSN 1548-7091. doi: 10.1038/nmeth.4292. URL http://www.nature.com/doifinder/10.1038/nmeth.4292.

[30] Allon Wagner, Aviv Regev, and Nir Yosef. Revealing the vectors of cellular identity with single-cell genomics. Nature Biotechnology, 34(11):1145–1160, nov 2016. ISSN 1087-0156. doi: 10.1038/nbt.3711. URL http://www.nature.com/articles/nbt.3711.

[31] F Alexander Wolf, Philipp Angerer, and Fabian J Theis. {{SCANPY}}: Large-Scale Single-Cell Gene Expression Data Analysis. 19:15. ISSN 1474-760X. doi: 10.1186/s13059-017-1382-0.

[32] Zonghan Wu, Shirui Pan, Fengwen Chen, Guodong Long, Chengqi Zhang, and S Yu Philip. A comprehensive survey on graph neural networks. IEEE transactions on neural networks and learning systems, 32(1):4–24, 2020.

[33] Jing Zhao, Xijiong Xie, Xin Xu, and Shiliang Sun. Multi-view learning overview: Recent progress and new challenges. Information Fusion, 38:43–54, nov 2017. ISSN 1566-2535. doi: 10.1016/J.INFFUS.2017.02.007.

