## appendix for "Multimodal single cell data integration challenge: results and lessons learned"

| Name | Affiliation |
|---|---|
| Carl P Walther | Baylor College of Medicine |
| Ramon Viñas | University of Cambridge |
| Brendan Panici | Baylor College of Medicine |
| Tamim Abdelaal | Leiden University Medical Center, Delft University of Technology, the Netherlands |
| Lieke Michielsen | Leiden University Medical Center, Delft University of Technology, the Netherlands |
| Ahmed Mahfouz | Leiden University Medical Center, Delft University of Technology, the Netherlands |
| Marcel JT Reinders | Delft University of Technology, Leiden University Medical Center, the Netherlands |
| Fei Wang | Weill Cornell Medicine |
| Zilong Bai | Weill Cornell Medicine |
| Hao Zhang | Weill Cornell Medicine |
| Matthew Brendel | Weill Cornell Medicine |
| Haotan Zhang | Weill Cornell Medicine |
| Nicholas Bartelo | Weill Cornell Medicine |
| Fenglei Fan | Weill Cornell Medicine |
| Ahmet Sarıgün | Middle East Technical University |
| Ahmet S Rifaioglu | Institute for Computational Biomedicine, Faculty of Medicine, Heidelberg University and Heidelberg University Hospital, Heidelberg, Germany; Department of Computer Engineering, İskenderun Technical University, Hatay, Turkey |
| Julio Saez-Rodriguez | Institute for Computational Biomedicine, Faculty of Medicine, Heidelberg University and Heidelberg University Hospital, Heidelberg, Germany |
| Deniz Germen | Middle East Technical University |
| Zoi Tsangalidou | Oxford University, Department of Statistics |
| Miriam Stricker | Oxford University, Department of Statistics |
| Arni Gunnarsson | Oxford University, Department of Statistics |
| Romain Fournier | Oxford University, Department of Statistics |
| Georgios Kalantzis | Oxford University, Department of Statistics |
| Hrushikesh Loya | Oxford University, Department of Statistics |
| Wanwen Zeng | Stanford University |
| Chencheng Xu | Tsinghua University |
| Tianyu Liu | University of Illinois at Urbana-Champaign |
| Grant Greenberg | University of Illinois at Urbana-Champaign |
| Ilan Shomorony | University of Illinois at Urbana-Champaign |
| Robert Lehmann | KAUST |

| | |
|---|---|
| Xabier Martinez De Morentin | Translational Bioinformatics Unit, Navarrabiomed, Complejo Hospitalario de Navarra (CHN), Universidad Pública de Navarra (UPNA), IdiSNA, Pamplona, Spain. |
| Minxing Pang | KAUST |
| David Gomez-Cabrero | KAUST |
| Narsis Kiani | Algorithmic Dynamics Lab, Department of Oncology-Pathology, & Center of Molecular Medicine, Karolinska Institutet, Stockholm, Sweden. |
| Jesper Tegner | KAUST |
| Wanli Wang | Baylor College of Medicine |
| Yu-Hsiu Chen | None |
| Sheng Wan | None |
| Chun-Hao Chao | None |
| Tung-Yu Wu | None |
| Ewa Szczurek | University of Warsaw |
| Marcin Możejko | University of Warsaw |
| Krzysztof Koras | University of Warsaw |
| Paulina Szymczak | University of Warsaw |
| Mike Phuycharoen | The University of Manchester |
| Chaozhong Liu | Baylor College of Medicine |
| Linhua Wang | Baylor College of Medicine |
| David Banh | AskExplain |
| Yanwan Dai | Baylor College of Medicine |
| Huidong Chen | Mass General Hospital/Harvard Medical School |
| Luca Pinello | Mass General Hospital/Harvard Medical School/BROAD |
| Jayoung Ryu | Harvard Medical School/Mass General Hospital |
| Justin Delano | Harvard Medical School/Mass General Hospital |
| Igor I | Novel Software Systems LLC |
| Xin-Ming Tu | University of Washington |
| Chen-Rui Xia | Peking University |
| Michaela A McCown | Baylor College of Medicine |
| Simon Steshin | Skolkovo Institute of Science and Technology |
| Alina Paronyan | Skolkovo Institute of Science and Technology |
| Rohan Gala | Allen Institute of Brain Science |
| Uygar Sümbül | Allen Institute of Brain Science |
| Roman Laddach | King's College London |
| Michael Shapiro | Francis Crick Institute |
| Sungyoon Kim | KAIST |
| HyeongJoo Hwang | KAIST |
| Seunghoon Hong | KAIST |
| Kee-Eung Kim | KAIST |
| Muhammad Irfan Khan | Turku University of Applied Sciences, Finland |
| Mohammad Ayyaz Azeem | Riphah International University, Pakistan |
| Najwa Laabid | University of Eatern Finland |
| Merja Heinäniemi | University of Eatern Finland |

| | |
|---|---|
| Janne Suhonen | University of Eatern Finland |
| Atanas Pavlov | SysMo Ltd. |
| Dean Palejev | Big Data for Smart Society (GATE) Institute, Sofia University, Bulgaria; Institute of Mathematics and Informatics, Bulgarian Academy of Sciences, Sofia, Bulgaria |
| Dimitar Jetchev | Inpher |
| Strahil Pastuhov | Nagoya University |
| Yuanfang Guan | University of Michigan, Department of Computational Medicine and Bioinformatics |
| Zhicheng Ji | Duke University |
| Changxin Wan | Duke University |

Table 1: NeurIPS 2021 Multimodal data integration competition participant authors

### Appendix B. Supplementary Figures

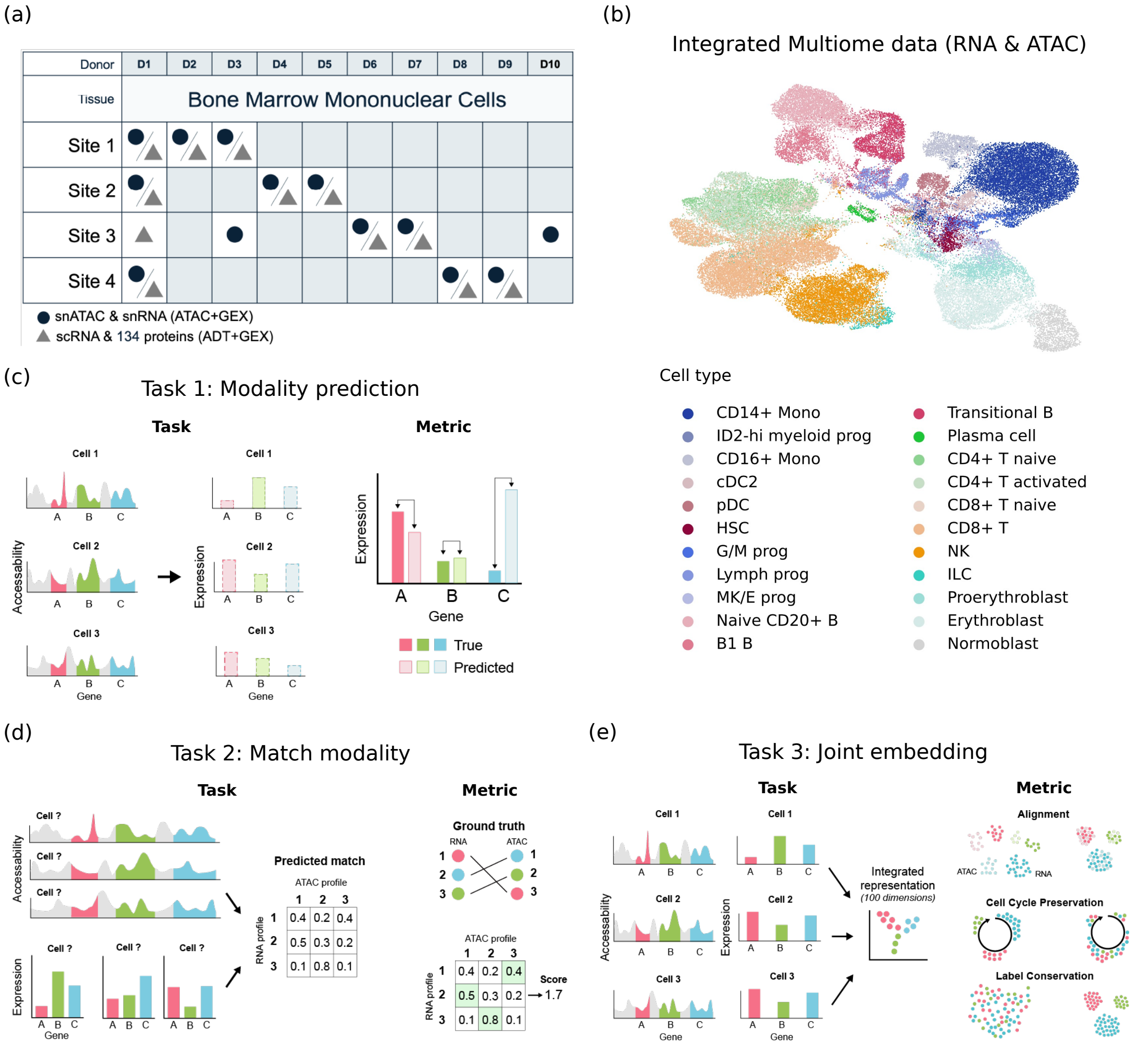


Figure 4: **Competition set-up.** **(a)** Experimental design to generate the multimodal single-cell benchmarking dataset with nested batch structure and **(b)** an example of an integrated representation of the Multiome data created using multiVI (Ashuach et al., 2021). **(c-e)** Conceptual figures of the three tasks and evaluation metrics used in the competition. Task 1 was the prediction of one modality from another based on paired training data and evaluated by RMSE. Task 2 was the matching of cells across modalities, evaluated by a match probability score. Task 3 was learning a joint embedding leveraging the variance of two modalities, evaluated by the removal of batch effects and conservation of biological signatures such as cell type and cell differentiation. See Luecken et al. (2021) for details.

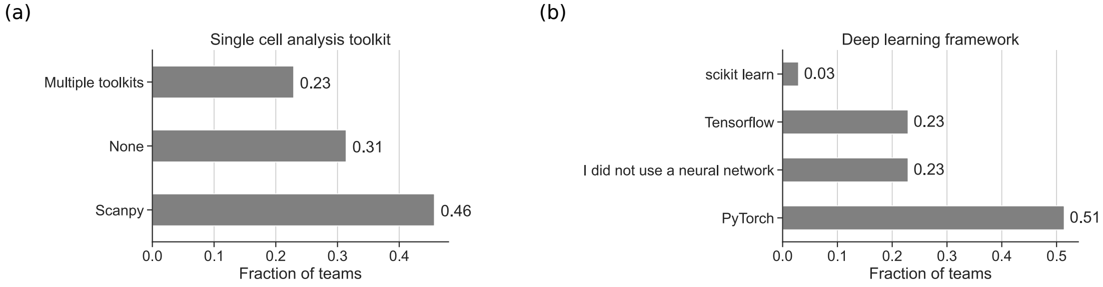


Figure 5: **Toolkit breakdown**. Software packages used by competitors for both single cell analysis **(a)** and deep learning **(b)** based on all survey responses.

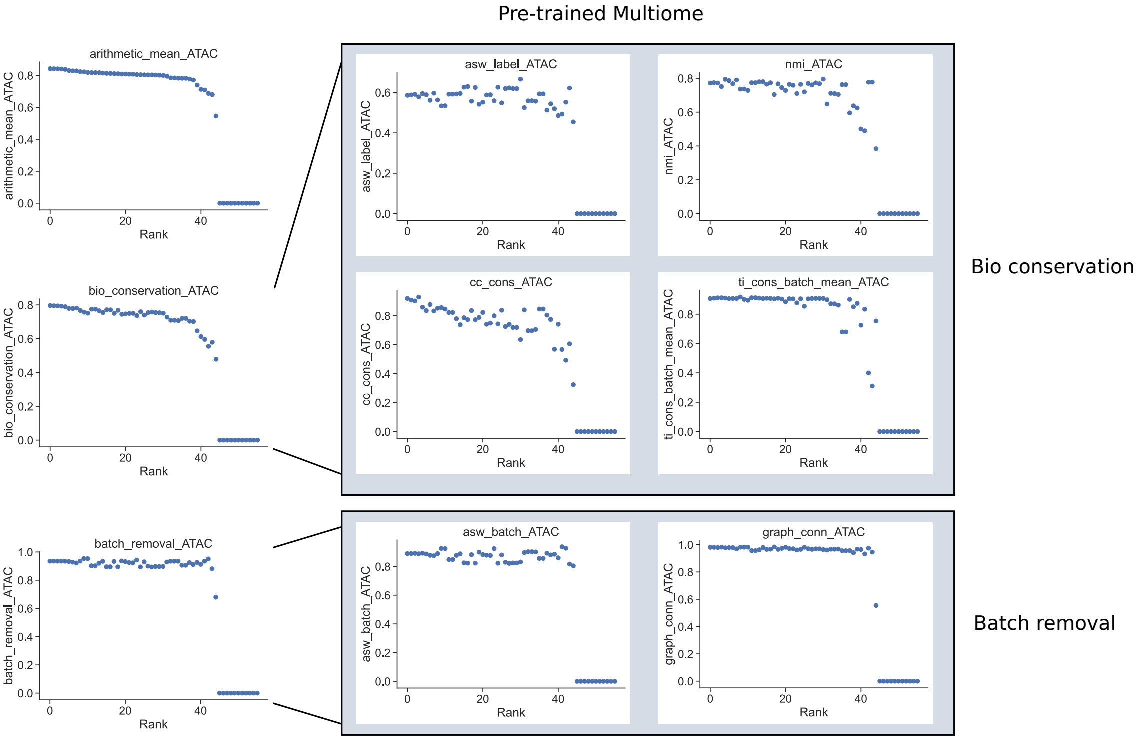


Figure 6: **Task3: joint embedding evaluation metrics by rank on the pre-trained Multiome subtask.** Top left shows the overall integration score, which is split up in batch removal and bio-conservation metrics. See Luecken et al. (2021) for detailed metric descriptions.

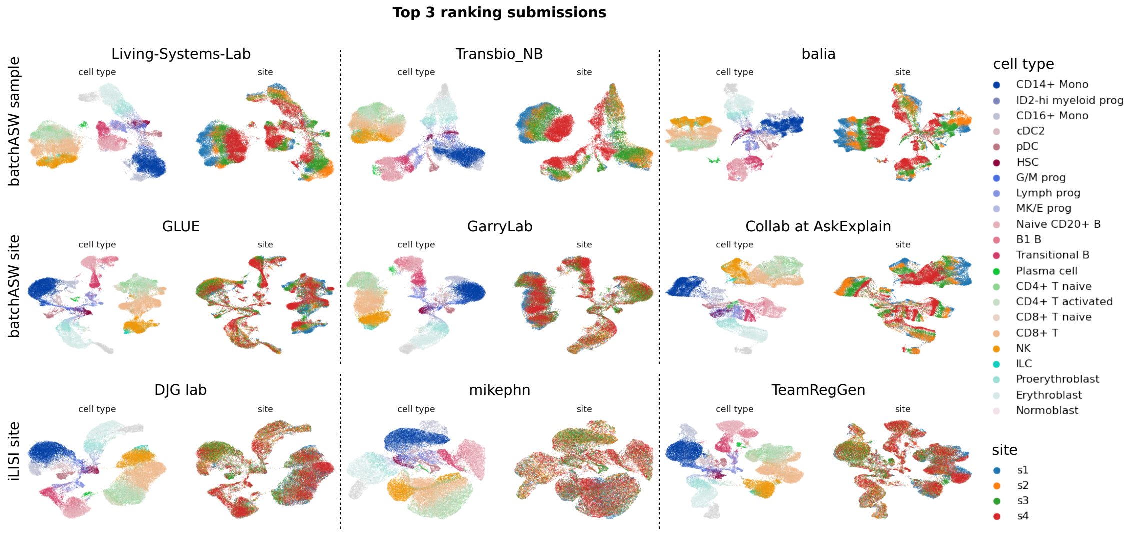


Figure 7: **Task 3: Extended joint embedding metric evaluation**. Additional best performing methods (left to right) ranked by batchASW sample, batchASW site and iLISI site.

#### Appendix C. Additional method descriptions

The code for all methods described here can be found at `https://github.com/openproblems-bio/neurips2021_multimodal_topmethods`.

##### C.1. Task 1: Modality prediction

**GEX→ATAC - 1st place: Living Systems Lab** Living Systems Lab used PCA to project the input and output modalities into 50 dimensions, and then trained a k-nearest neighbors regression model between these representations using 25 neighbors with the Minkowski distance metric (Fig 8). The full-dimensional output was reconstructed using the saved principal axes. The dataset was split into five folds stratified by the batches. The model was trained on each fold, and the outputs were then averaged. Hyperparameters were optimized in the local validation and public testing sets using the RMSE metric.

**ATAC→GEX - 1st place: Cajal** Cajal is a deep neural network method for the prediction of one modality from another. Cell-type specific features are selected as input, based on differential expression (Wilcox test) or differential accessibility (T-test after binarization of data), using an annotated reference dataset. The analysis of this reference dataset is performed using SCANPY (Wolf et al.). Feature selection is not performed when the input modality is ADT, due to the small number of proteins profiled in CITE-seq datasets. Total counts, median total counts per batch and the standard deviation in total counts per batch are also calculated and used as input features. Values for the selected features are centered and scaled before being input into the neural network. The network architecture consists of

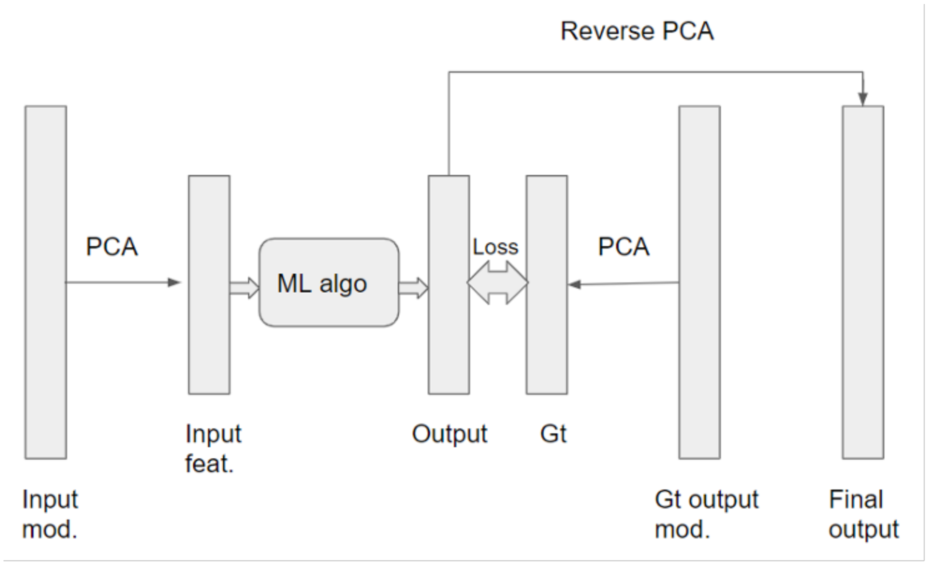


Figure 8: **GEX→ATAC: Living Systems Lab**.

a dropout layer, 3 to 6 hidden layers which use the ReLU activation function, an output layer with no activation function (linear regression), and a final Lambda layer. This Lambda layer serves to clip the output to biologically reasonable values (the range present in the training data). The neural networks are implemented using TensorFlow (Abadi et al., 2016) and, during the training and validation phases, KerasTuner (O'Malley et al., 2019) was used to optimize hyperparameters (percentage dropout, number of hidden layers, number of nodes per layer).

Team Cajal initially sought to create a heteroencoder, however discovered that implementing a bottleneck architecture did not result in optimum performance across tasks. Thus, the hyperparameter search was restricted to $10-90\%$ dropout, 3 - 6 hidden layers (after initial experimentation showing less than 3 resulted in reduced performance), and $10-800$ nodes per layer (range chosen loosely based upon single-cell deep learning tasks reported in the literature). A number of approaches were attempted that did not improve performance, including pre-training as an autoencoder, L2 regularization, one-hot encoding of cell type as inferred by label transfer from the challenge reference dataset using ingest, and the use of additional datasets during training. To prevent overfitting, Team Cajal found that early stopping was necessary before the addition of a dropout layer. After this addition, early stopping was no longer essential, although no improvement in performance on the validation dataset was seen after 50 epochs. The final model parameters are detailed in Table 2.

**GEX→ADT - 1st place: Guanlab - dengkw** Team Guanlab - dengkw built a Kernel Ridge Regression (KRR) model with the Radial Basis Function (RBF) kernel $k(x_i, x_j) = exp(-\frac{|x_i-x_j|^2}{2l^2})$. Here, the $d(.,.)$ is the Euclidean distance (Fig 9). Processed data was used as input to the model. Data processing was done by: (1) concatenating the data from modality 1 (Mod 1) in the training and test set with an outer join by the features and labeling them

| Modality | Dropout | Layer Sizes |
|---|---|---|
| ATAC - GEX | 0.8 | 320, 360, 620, 440 |
| GEX - ATAC | 0.8 | 500, 20, 110, 430, 310 |
| GEX - ADT | 0.2 | 170, 300. 480, 330, 770 |
| ADT - GEX | 0.2 | 180, 140, 520 |

Table 2: Cajal modality prediction submission parameters

with ”train” and ”test”; (2) applying a truncated SVD to the combined matrix for dimension reduction; (3) applying a truncated SVD to the training data from modality 2 (Mod 2); (4) applying row-wise z-score normalization on the reduced matrix and splitting the matrix by the labels. Finally, the KRR model was fit with the normalized training matrix and the processed Mod 2 data. Outputs from predicting the test matrix were mapped back to the dimensions of test Mod 2 data by multiplying by the right singular matrix.

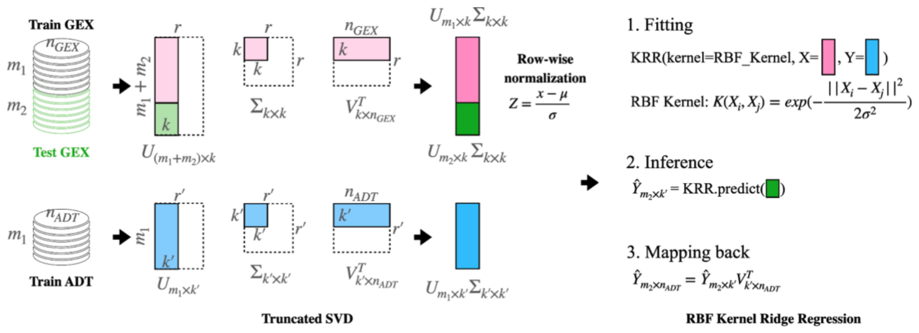


Figure 9: **GEX→ADT: Guanlab - dengkw**. Model overview.

In the final submission, in order to overcome memory limitations, Team Guanlab - dengkw applied a training strategy consisting of random sampling and assembling (Fig 10). First, batches were randomly shuffled in the training data and two models were trained on the first and second half batches of the data. Then the steps were repeated 5 times and 10 models were generated with 10 outputs. The final predictions were the average of these outputs.

There are 4 tunable parameters in the model: the number of components for Mod 1 and Mod 2, the length scale $l$ in the RBF kernel, and the regularization strength $\alpha$ in ridge regression. These parameters were determined by cross-validation performances on the Phase I data. They are listed in Table 3. For the ADT2GEX task no feature reduction was applied to the ADT data.

**ADT→GEX - 1st place: Novel** Team Novel used an MLP with encoder-decoder structure, with most of the model capacity contained in the decoder given the high dimensional output. Although ATAC and GEX modalities were compressed via latent semantic indexing (LSI) for other input modalities, the ADT input data was not pre-processed due to its already low dimensionality. To avoid overfitting, the model was regularized using dropout

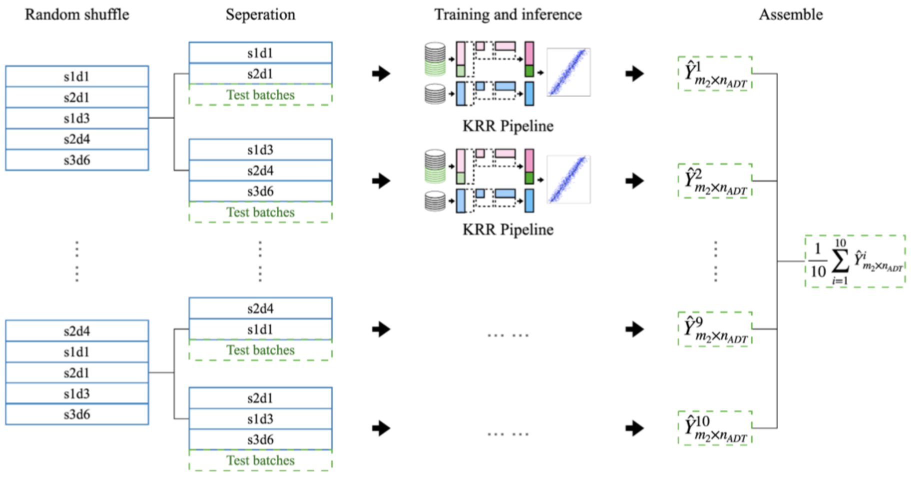


Figure 10: **GEX→ADT: Guanlab - dengkw**. Strategy to overcome memory limitations.

| **Task** | **# of Mod 1 components** | **# of Mod 2 components** | $l$ | $\alpha$ |
|---|---|---|---|---|
| GEX2ADT | 300 | 70 | 10 | 0.2 |
| ADT2GEX | None | 50 | 10 | 0.2 |
| GEX2ATAC | 1000 | 50 | 10 | 0.1 |
| ATAC2GEX | 100 | 50 | 10 | 0.1 |

Table 3: Guanlab - dengkw modality prediction submission parameters

before every hidden layer, as well as and weight decay. Alternative regularization techniques that were explored include batch swap noise and a variational prior on the latent variables, however these did not lead to beneficial results. Notably, in some cases a higher test metric score was achieved after training on a smaller data set rather than the full data set. For example, using the phase 1 data as a training dataset proved best to train the ADT2GEX model. The optuna framework was used to tune hyperparameters (LSI dimension, dropout rate before every hidden layer, depth, layer widths, activation function, learning rate, optimizer and weight decay). The optimal hyperparameter values are detailed in Table 4.

**ADT→GEX - 2nd place: Living Systems Lab** Living Systems lab used a similar pipeline for the ADT2GEX subtask as for the winning submission in the GEX2ATAC task (see above). They trained a neural network with 2 hidden layers, a ResNet-style neural network with skip connections and a batch prediction head with 3 hidden layers; and catboost models with Bernoulli bootstrapping and Bayesian bootstrapping on 5 folds. These models were ensembled by averaging the output features.

**Overall: DANCE** Team DANCE leverages a bipartite graph between cells and different modalities (GEX, ADT, and ATAC) to represent interactions. They utilize graph neural networks adapted from Battaglia et al. (2018); Kipf and Welling (2016) and Wu et al.

| | **GEX2ATAC** | **ATAC2GEX** | **ADT2GEX** | **GEX2ADT** |
|---|---|---|---|---|
| # HL | 3 | 3 | 3 | 3 |
| HL dims | [1024, 256, 2048] | [2048, 2048, 512] | [512, 512, 512] | [512, 512, 2048] |
| HL dropout rates | [0.30, 0.11, 0.14] | [0.26, 0.18, 0.25] | [0.0, 0.0, 0.0] | [0.20, 0.15, 0.17] |
| Activation functions | GeLU | GeLU | GeLU | GeLU |
| Learning rate | 1e-05 | 8e-05 | 4e-04 | 3e-05 |
| Optimizer | AdamW | AdamW | Adam | AdamW |
| Batch size | 64 | 256 | 64 | 32 |
| Weight decay | 4e-04 | 7e-04 | 1e-05 | 1e-02 |
| # LSI components | 256 | 256 | - | 256 |

Table 4: Novel modality prediction submission parameters; HL - hidden layers; dims - dimensions

(2020) to exploit the structural information of this bipartite graph. Specifically, the solution consists of three major components: (1) bipartite graph construction, (2) heterogeneous graph convolution, and (3) representation concatenation. In the first step, a bipartite graph is constructed between modality (GEX, ADT and ATAC) and cell nodes. In the GEX2ADT and GEX2ATAC tasks, they additionally introduced pathway data from hallmark gene sets (i.e. sets of related genes). Based on the original bipartite graph, gene features which are shown to be related in the pathway dataset were linked.

The initial node embedding of feature nodes is a one-hot index of each feature, thus, the node embedding matrix for all feature nodes $\hat{X}_{feat} = R^{k \times k}$ , is an identity matrix (a.k.a $\hat{X}_{feat} = I_k$ ), where $k$ is the number of modality 1 (Mod1) features. To model the batch effect, initial cell embeddings were computed from batch features, the statistical features of all the cells in one batch. Eventually, for each cell a 9-dimensional batch feature was obtained, so that the batch features for all cell nodes could be described as $\hat{X}_{batch} \in R^{N \times 9}$.

This graphs was used as the basis for heterogeneous graph convolution. In the heterogeneous graph convolution, parameters for different types of edges were separated. In each convolution layer three sets of aggregation approaches were simultaneously completed to aggregate the information from different types of neighbors to the target node. The node embeddings were then updated with residual connection and a weighted sum of aggregation results:

$$X_{feat}^{l} = X_{feat}^{l-1} + \alpha * AGG_{c2f} + (1 - \alpha) * AGG_{f2f} \quad (1)$$

$$X_{cell}^{l} = X_{cell}^{l-1} + AGG_{f2c} \quad (2)$$

where $l$ is the number of current layer, $AGG_{f2f}$ is the pathway aggregation (i.e., the aggregation from feature to feature), $AGG_{c2f}$ is the cell-feature aggregation, $\alpha$ is a scalar hyper-parameter to control the ratio between those two aggregations, and $AGG_{f2c}$ is the aggregation from features to cells. Finally, Team DANCE took the embeddings of cell nodes from each convolution layer, concatenated them and projected them to the space of downstream task via a linear transformation.

##### C.2. Task 2: Match modality

**All subtasks - 2nd place: Novel** Team Novel approached the match modality task using a metric learning approach and post-processing adapted from graph theory. For each data type, a shallow MLP encoder was used to learn spherical sample embeddings. ATAC and GEX input data was preprocessed by latent semantic indexing (LSI) transformation (a TF-IDF transformation followed by SVD). In contrast to their approach in the Predict Modality task, the first LSI component was discarded as it was broadly associated with technical variation. ADT input data was directly fed into the network due to its lower dimensionality. The encoders were trained by minimizing the symmetric cross-entropy of cosine similarities between sample pairs. This approach was inspired by CLIP, a model designed to connect text and image data (Radford et al., 2021). Matching probabilities were inferred in a two step procedure: (1) evaluation of a dense pairwise cosine similarity matrix based on the embeddings of each modality; and (2) finding a maximum weight matching in a bipartite graph, where each vertex represents a sample in the data. To ensure computational feasibility, the input matrix was sparsified by discarding elements smaller than their corresponding row- and column-wise 0.995 quantiles. This way, the matrix was greatly sparsified while retaining values in all rows and columns. The proposed algorithm for sample matching is making hard bets rather than soft probability assignments. It is intrinsically symmetric so its output and metrics are consistent for both directions of the matching problem (i.e. ADT2GEX and GEX2ADT). For the ATAC2GEX and ADT2GEX subtasks the model was trained on Phase 2 and Phase 1 data, respectively, for 7000 epochs. Due to the high cost of the bipartite matching procedure, the competition metric was evaluated on each epoch using probabilities obtained by a row- and column-wise softmax averaged for the modality pair. The hyperparameters for the embedding model were found by an extensive search using the optuna framework and are shown in Table 5.

##### C.3. Task 3: Joint embedding

**Multiome, pre-trained and CITE, pre-trained: Amateur** Team Amateur's method "joint embedding with a regularized autoencoder" (JAE) was inspired from previous work on scDEC (Liu et al., 2021), which aims at simultaneous deep generative modeling and clustering of single-cell data. Here, the scDEC model was simplified by removing the discriminator networks and adding constraints to the encoder latent space so that JAE requires latent features to recover more biological knowledge, including cell type, batch, and cell-cycle phase.

In the JAE model, each modality (except ADT) is first SVD transformed (e.g. to 100 components) and concatenated. The information from cell annotations (e.g., cell label, cell cycle score, and cell batch) is incorporated to constrain the structure of latent features. In this manner some latent features should recover the cell type information and some should recover the cell cycle score. Batch-related features should recover batch labels as randomly as possible to potentially eliminate this effect. Some features in the latent space were left without constraint to ensure the flexibility of network. JAE was pre-trained using the provided annotated datasets in an end-to-end fashion (Adam optimizer, $lr = 10^{-4}$), where multiple loss functions were used, including autoencoder reconstruction loss, cell type prediction cross entropy loss, cell cycle phase score mean squared error (MSE) loss, and batch

| | **ADT ↔ GEX** | **ATAC ↔ GEX** |
|---|---|---|
| Learning rate | 8e-05 | 6e-04 |
| Optimizer | AdamW | AdamW |
| Weight decay | 0 | 0 |
| Batch size | 2048 | 16384 |
| Embedding dimension | 64 | 256 |
| # HL in GEX encoder | 2 | 2 |
| # HL in Mod 2 encoder | 2 | 1 |
| HL dims (GEX) | [1024,512] | [1024,1024] |
| HL dims (Mod 2) | [512, 2048] | [2048] |
| HL dropout rates(GEX) | [0.01, 0.25] | [0.54, 0.4] |
| HL dropout rates(Mod 2) | [0.02, 0.3] | [0.66] |
| Initial temperature (symmetrical cross-entropy [log]) | 3.46 | 3.06 |
| Number of LSI components | 128 for GEX | 512 for ATAC, 64 for GEX |

Table 5: Novel match modality submission parameters; HL - hidden layers; dims - dimensions

loss. These losses were balanced, resulting in a total loss of $0.7\,\mathrm{AE} + 0.2\,\mathrm{CT} + 0.05\,\mathrm{CC} + 0.05\,\mathrm{Batch}$. In the online testing stage where cell annotations were not available, the model was fine-tuned only based on the autoencoder reconstruction loss. This was done for 1 epoch with learning rate of $10^{-4}$ and 2 epochs with a learning rate of $2 * 10^{-5}$ for the Multiome and CITE-seq data, respectively. Finally, all the features in the JAE latent space were used as the joint embedding.

**Multiome, online: Living Systems Lab** The Living Systems Lab team solved this task using a concatenated autoencoder (cAE) architecture. The cAE consists of two encoders, each with a single hidden layer that is concatenated and then projected down to an embedding layer, a concatenated inverse layer, and two decoder layers. Before training the cAE, the data were pre-processed by (1) identifying highly variable genes and (2) setting a threshold to filter peaks that are rarely identified (3% or less of cells). Furthermore, a weight orthogonality constraint was applied to both encoding layers, enhancing the embedding's discriminability and representability.

Each encoder, which receives data from a different modality, has 64 dimensions, leading to 128 dimensions after concatenation. The embedded layer, which contains the two domain's integrated low-dimensional representation, again consists of 64 dimensions. ReLU was used as an activation function for all layers with a dropout value of 0.1 applied to two encoding layers, except in the bottleneck layer, where a linear activation function was used. Training was performed with the ADAM optimizer for 600 epochs at a learning rate of $lr = 0.0001$ and a batch size of 32. An MSE loss function was used.

**CITE, online: Guanlab - dengkw** Team Guanlab - dengkw solved the CITE online subtask using the same linear approach as in task 1. Truncated SVD was applied to the

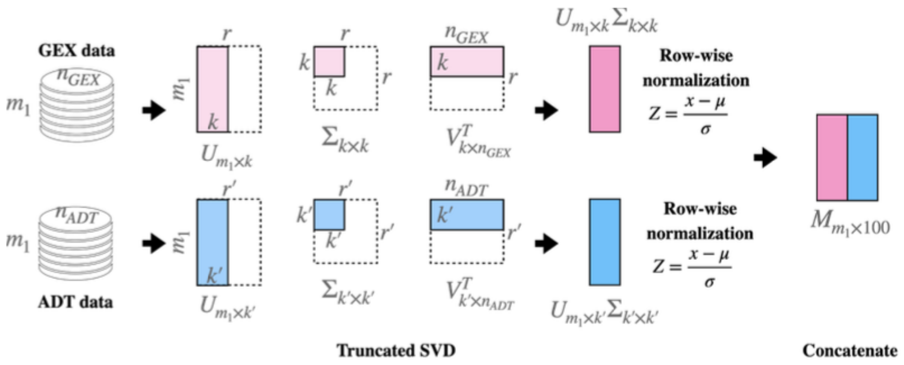


Figure 11: **Joint embedding (CITE, online): Guanlab - dengkw**.

modality 1 and the modality 2 data, respectively, and normalized on the rows for each matrix. The output is a simple concatenation of the reduced features. The numbers of components for each modality were determined by submission performance in experiments using the Phase 1 data. In the task of embedding the CITE data, the dimensions for GEX and ADT are 73 and 27, respectively. More details and a schematic diagram can be found in Figure 11.
